# Hybrid modeling framework for bioprocesses with minimal prior knowledge and limited data

**DOI:** 10.1101/2025.11.14.688550

**Authors:** Carlos Martínez, Facundo Rocha Calvette, Marielle Péré, Mauricio Barrientos, Sebastián Ossandón

**Affiliations:** Escuela de Ingeniería Bioquímica, Pontificia Universidad Católica de Valparaíso, Av. Brasil 2085, 2362803 Valparaíso, Chile; Aix Marseille Univ, INSERM, MMG, Marseille, France; Université Côte d’Azur, Institut de Pharmacologie Moléculaire et Cellulaire (IPMC, CNRS UMR 7275, Inserm 1323), 06560 Valbonne Sophia Antipolis, France; Institut Hospitalo-Universitaire (IHU) RespirERA, Nice, France; Université Côte d’Azur, Inria, INRAE, CNRS MACBES Team, 06902 Valbonne Sophia Antipolis, France; Instituto de Matemáticas, Pontificia Universidad Católica de Valparaíso, Blanco Viel 596, Cerro Barón, Valparaíso, Chile

**Keywords:** Data-driven modeling, Batch culture, Neural network, Parameter estimation, Small datasets

## Abstract

Hybrid models that couple mechanistic ordinary differential equations (ODEs) with neural networks are increasingly used in bioprocess engineering, yet most published approaches assume either substantial prior knowledge or relatively large datasets. This work proposes a hybrid modeling framework for early-stage bioprocess development, where only a few batch experiments are available and standard artificial intelligence (AI) techniques are difficult to apply. The mechanistic structure is constructed using only qualitative, widely accepted biological constraints (e.g., non-negativity, zero-invariance, and biomass-mediated interactions), while unknown functional dependencies are learned by a feedforward neural network embedded in the ODE right-hand side. To exploit the natural organization of batch data, we introduce a minibatch training strategy in which each minibatch corresponds to one entire batch experiment, combined with regularization to mitigate overfitting. We demonstrate the approach on (i) synthetic *Escherichia coli* growth with overflow metabolism and (ii) experimental astaxanthin production by *Xanthophyllomyces dendrorhous*. In both cases, models trained from as few as three batch experiments accurately predict an unseen validation batch and the learned neural components recover biologically consistent patterns. Thus, the framework contributes to AI by enabling constrained neural differential models that learn interpretable dynamics from limited, structured data, with applications to early-stage bioprocess engineering.

## 1 Introduction

The concept of hybrid modeling for bioengineering was first proposed by [21] for fermentation process control and has been further developed since the 2000s [7, 17]. Hybrid models integrate a data-driven and a parametric model (mechanistic/empirical component) into a unified system that benefits from both physical properties and data-driven aspects. The mechanistic part is generally described by a system of ordinary differential equations (ODEs), and the data-driven component is described by a neural network (NN).

Hybrid models have been successfully applied to represent several bioprocesses [1], including the production of biopharmaceuticals with mammalian cells [20, 16], the production of recombinant proteins with Escherichia coli [29, 3], and the production of amoxicillin by biocatalysis [26]. However, most published studies rely on relatively large datasets to train hybrid models [29, 16, 19, 22]). For example, [16] reported results based on 81 time-series datasets obtained experimentally, while [18] used 25 synthetic time-series datasets derived from model simulations. However, in many practical situations, particularly during early-stage process development, only limited experimental data are available, making model training difficult and prone to overfitting. At the same time, this is also the stage where a model could be most useful: supporting the planning of future experiments, narrowing the operational space, and reducing the cost of empirical trial-and-error.

In addition to data scarcity at early development stages, constructing hybrid models presents challenges of its own. Incorporating uncertain or inaccurate prior mechanistic knowledge can degrade predictive performance and lead to ill-conditioned estimation problems [23]. This issue is particularly pronounced in the early research phases when reliable mechanistic knowledge is often limited. Additionally, the introduction of unknown parameters into the mechanistic component further complicates model training, requiring the simultaneous estimation of mechanistic and data-driven parameters, a task known to be computationally demanding and prone to identifiability issues [25]. To mitigate these difficulties, two-step training strategies have been proposed, in which mechanistic and data-driven components are fitted separately before integration [31]. However, decoupling the estimation steps does not guaranty global optimality of the hybrid model.

The objective of this paper is to develop a hybrid modeling framework that relies on minimal prior knowledge and only a few datasets. We focus on data obtained from batch cultivations, a widely used setup in early-stage bioprocess research [30, 4, 12, 8]. In batch mode, the culture evolves in a closed system without fresh medium addition, producing a single time-series trajectory per run that typically includes measurements of biomass and substrate or metabolite concentrations. Batch runs are routinely repeated under different conditions, providing a sparse yet informative basis for model inference. By a model with minimal prior knowledge, we refer to a formulation that incorporates only general, universally accepted physical and biological constraints (e.g., non-negativity, no spontaneous generation of biomass, and biomass-mediated reactions). This keeps the model independent of explicit kinetic assumptions while still ensuring physical plausibility and generalization.

Our framework addresses both the construction of the hybrid model and the use of data during training. On the modeling side, we impose fundamental biological constraints. By relying only on well-established principles, we avoid introducing uncertain assumptions that could degrade model performance. On the training side, we introduce a minibatch training scheme based on the ADAM optimizer [11], in which each minibatch corresponds to the data associated with a single batch experiment. At each iteration, the optimization algorithm refines the model using data from one batch experiment, mimicking the sequential process of calibration across experiments. This approach differs from conventional methods [18], where randomly sampled data points form minibatches. Such conventional strategies are less natural in the context of batch bioreactors, where data are inherently organized by experiments (“minibatch”) and may require repeated ODE integrations across multiple initial conditions in each optimization iteration. To avoid overfitting, regularization techniques such as early stopping, dropout, or parameter penalization can be applied [15]. In particular, dropout regularization [27], originally developed for deep neural networks, has recently been implemented in hybrid modeling frameworks [18, 22]. Dropout works by randomly deactivating a fraction of neurons during training, preventing the network from relying too heavily on specific connections. As stated by [18], the stochastic nature of dropout makes the model not only less prone to overfitting but also less dependent on the initial configuration of parameters. Thus, we incorporate dropout regularization into the training algorithm to reduce overfitting.

The paper is organized as follows. Section 2 presents the proposed modeling framework. We begin by introducing a general hybrid model that serves as a flexible foundation, and then we describe the procedure to adapt this model for specific bioprocess scenarios. This section also details the algorithm developed to calibrate hybrid models within the proposed framework. Section 3 demonstrates the approach through two case studies: the first employs a synthetic dataset for E. coli, while the second applies the method to experimental data on astaxanthin production by *X. dendrorhous*. Finally, the paper concludes with a summary of the main findings and a discussion of perspectives for future research.

## 2 Hybrid modeling framework

### 2.1 Hybrid model

Consider a closed system in which microorganisms grow. Let

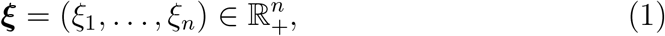

denote the state vector of the process, where ℝ_+_ represents the set of nonnegative real numbers, and let ℐ := {1, …, *n* } be the corresponding index set. In typical bioprocess applications, the components of ***ξ*** represent the concentrations of biomass, substrates, and metabolic products. We propose the following set of differential equations to describe the dynamics of ***ξ***:

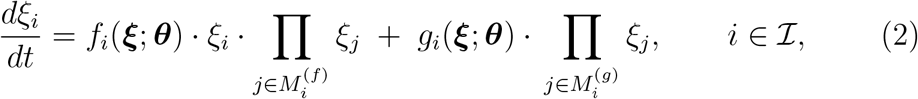

where *f*_*i*_ : ℝ^*n*^ × Θ →ℝ and *g*_*i*_ : ℝ^*n*^ × Θ →ℝ_+_ are described by feedforward neural networks (FNN). An FNN is a standard machine-learning model used to approximate unknown nonlinear relationships directly from data (details are presented in Section 2.3). The set Θ denotes the parameter space of the FNN. A specific configuration of these parameters is represented by ***θ*** ∈ Θ. Note that the functions *g*_*i*_, *i* = 1, …, *n* take values in ℝ_+_, ensuring non-negative outputs. The index sets 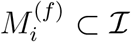 and 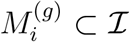 specify state variables that multiply the functions *f*_*i*_ and *g*_*i*_, respectively. The particular choice of these sets depends on the system under study and reflects qualitative knowledge, such as biomass-mediated uptake or product formation. By convention, an empty product equals one, *i*.*e*. Π_*j*∈∅_ *ξ*_*j*_ := 1; thus, if 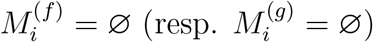.

For each state variable *ξ*_*i*_, the right hand side (RHS) in (2) involves two functions, *f*_*i*_ and *g*_*i*_. The function *g*_*i*_ represents the generation of the metabolite *ξ*_*i*_ that can occur independently of its current concentration. The function *f*_*i*_ captures the part of the dynamics that vanishes in the absence of *ξ*_*i*_; this follows from observing that *f*_*i*_ is multiplied by *ξ*_*i*_. The products of the variables that multiply each function indicate which state variables are required to mediate changes in other state variables.

The following proposition states that the solutions to (2) are non-negative, provided that the initial conditions are also non-negative.

#### Proposition 1

**(Non-negative state variables)** *If ξ*_*i*_(0) ≥ 0 *for all i* ∈ℐ, *then ξ*_*i*_(*t*) ≥ 0 *for all t* ≥ 0 *and all i* ∈ ℐ.

**Proof**. By contradiction, let us assume that there is a time *t* such that 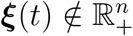. This implies the existence of a time *t*^*′*^ *>* 0 such that 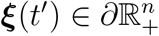 (boundary of the positive cone) and an index *i*^*′*^ ∈ ℐ such that

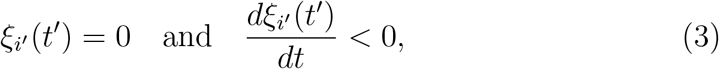

Otherwise, *ξ* cannot cross the boundary of 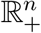. Since 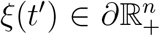, we have that *ξ*_*i*_(*t*^*′*^) ≥ 0 for all *i* ∈ ℐ. This implies that

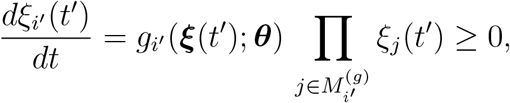

which contradicts (3). This completes the proof. □

The following proposition shows the flexibility of the formulation (2) in describing different scenarios in which one state variable mediates the dynamics of other variables.

#### Proposition 2

*Let i* ∈ *I. Then:*

a. *If g*_*i*_ ≡ 0, *then in the absence of ξ*_*i*_ *there is no net change in ξ*_*i*_.
b. *If* 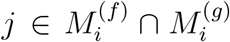 *(i*.*e*., *ξ*_*j*_ *appears as a multiplicative factor on the RHS of the equation for ξ*_*i*_*), then in the absence of ξ*_*j*_ *there is no net change in ξ*_*i*_.
c. *If* 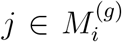 *(i*.*e*., *ξ*_*j*_ *multiplies g*_*i*_ *on the RHS of the equation for ξ*_*i*_*), then when both ξ*_*j*_ *(e*.*g*., *biomass) and ξ*_*i*_ *(e*.*g*., *product) are absent, there is no net change in ξ*_*i*_.

**Proof**. The proof follows directly from noting that, in each situation, the derivative of the state variable becomes zero under the given conditions. □

Based on Proposition 2, we propose a systematic procedure for constructing models that follow the structure of (2) and are tailored to the specific system under study. The procedure is structured into three steps, which are applied to each state variable individually.

### 2.2 Modeling procedure

#### Procedure 1 (Construction of the Hybrid Model)

*The construction of the hybrid model proceeds through three steps, which progressively transform a first formulation into a model suited to the system under study. Let us consider the state vector* ***ξ*** *from (1)*.

#### Step 1: Initialization

*Write the following ODE system, hereafter referred to as the* seed model:

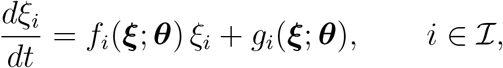

*where f*_*i*_ *and g*_*i*_ *correspond to the data-driven components of the model. This seed model corresponds to (2) when* 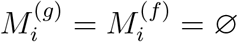.

#### Step 2: Enforcing self-invariance at zero

*For each state variable ξ*_*i*_, *assess whether it can emerge in the absence of itself. If spontaneous appearance is not possible (e*.*g*., *for biomass or substrate in closed systems), the generation term is removed by setting*

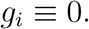

#### Step 3: Introducing mediated interactions

*Determine which state variables are required for any variation or formation of ξ*_*i*_. *Variables that modulate all changes of ξ*_*i*_ *multiply the entire RHS, while those that mediate its generation when ξ*_*i*_ = 0 *(for instance, biomass enabling product synthesis) multiply only the corresponding generation term*.

We refer the reader to Section 3 to see examples of how Procedure 1 leads to the construction of models.

##### Remark

Unlike classical bioprocess hybrid models, which are typically derived from mass balances [17, 20, 16, 18], the approach adopted here does not rely on any explicit conservation equations or mechanistic rate laws. Instead, the model is constructed from a set of qualitative constraints that encode essential biological principles, such as zero-invariance, biomass-mediated interactions, and the impossibility of spontaneous formation of certain states. These constraints define the admissible structure of the RHS of the ODEs, while the unknown functional dependencies are learned directly from data through neural networks. This strategy enables the formulation of dynamic models that remain biologically consistent without requiring detailed prior knowledge of stoichiometry, reaction pathways, or kinetic forms.

##### Remark

Note that physical or operational components, such as gas–liquid mass transfer or feed and dilution rate terms, can be readily incorporated into (2) to describe open systems such as continuous or fed-batch reactors. However, if operational parameters affect the kinetics, then they should be considered in the input layer of the neural network.

### 2.3 Neural network

The functions *f*_*i*_(***ξ***; ***θ***) and *g*_*i*_(***ξ***; ***θ***) in (2) are parameterized by an FNN with *L* layers. The input layer, denoted by **a**^(0)^, corresponds to the state variables; that is, **a**^(0)^ = ***ξ***. The subsequent hidden layers are computed iteratively as

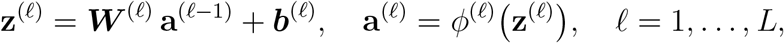

where *ϕ*^(*ℓ*)^ denotes the activation function at layer *ℓ*. For the hidden layers (*ℓ* = 1, …, *L* − 1), the activation function is chosen as *ϕ*^(*ℓ*)^(*u*) = tanh(*u*). The output layer produces

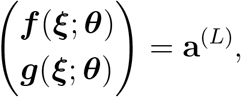

where ***f*** = (*f*_1_, …, *f*_*n*_)^*T*^ and ***g*** = (*g*_1_, …, *g*_*n*_)^*T*^. The activation function at the output layer is defined elementwise as *ϕ*^(*L*)^(*u*) = *u* for components corresponding to *f*_*i*_, and *ϕ*^(*L*)^(*u*) = ReLU(*u*) for components corresponding to *g*_*i*_. This choice ensures that all *g*_*i*_ functions remain non-negative while allowing *f*_*i*_ to take any real value. Finally, the full set of fitting parameters associated with the FNN is given by ***θ*** = {***W*** ^(*ℓ*)^, ***b***^(*ℓ*)^}_*ℓ*=1,…,*L*_.

### 2.4 Available data

In a typical experimental setup aimed at characterizing microbial kinetics, several independent batch experiments are conducted under varying conditions [30, 4, 12, 8]. Let *m* denote the total number of batch experiments. We will assume that each experiment provides a time series of measurements for all the state variables. Specifically, for the *j*-th experiment, the available data are of the form,

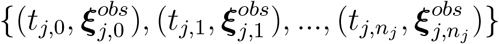

where *t*_*j,k*_ denotes the *k*-th observation time, and 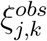 represents the corresponding measured concentrations of the state vector. We assume that all state variables are observed; however, in practice, some variables may be unmeasured or only indirectly inferred. The proposed framework can be readily extended to such cases without loss of generality.

In machine learning, it is standard practice to divide the available data into training and validation sets. Here, this division is performed at the *experiment level*. We introduce two index sets, *J*_train_, *J*_val_ ⊆ {1, …, *m*}, corresponding to the experiments used for training and validation, respectively. Training experiments are used to fit the neural-network parameters within the hybrid model, while validation experiments are held out to assess generalization and detect overfitting. A third set *J*_test_ may also be included to evaluate predictive performance on completely unseen conditions.

### 2.5 Training algorithm and implementation

We define the following loss function for the *j*-th batch experiment:

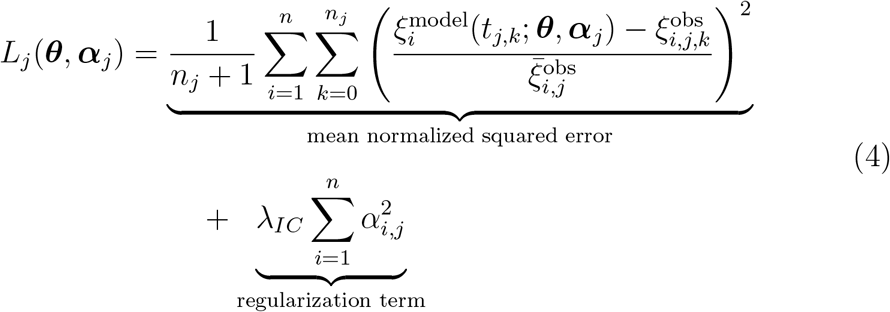

where *n*_*j*_+1 is the number of observation points in experiment *j*, 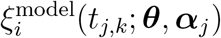 is the model-predicted value of state *i* at time *t*_*j,k*_, using parameters ***θ*** and ***α***_*j*_, 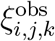 is the corresponding measurement, and 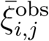 denotes the mean observed value of variable *i* in experiment *j*, used to normalize residuals. The vector of parameters ***α***_*j*_ := (*α*_1,*j*_, …, *α*_*n,j*_), which must be estimated, collects the parameters that define the initial condition of experiment *j*. Specifically, the model uses

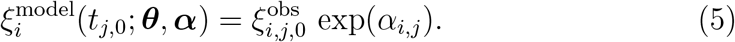

In this way, *α*_*i,j*_ = 0 recovers the measured initial value and nonzero values allow proportional adjustments while preserving non-negativity. The regularization term in (4) penalizes large deviations of the learned initial conditions from the measured ones, helping control the trade-off between fitting accuracy and robustness.

The global training loss is obtained by aggregating the contributions of all training experiments. We define it as the average of the individual losses:

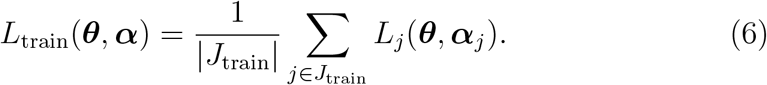

We use | *J*_train_ | to denote the cardinality of the index set *J*_train_. For validation experiments, the initial conditions are not estimated but directly taken from the observed data. Accordingly, the validation loss is defined as

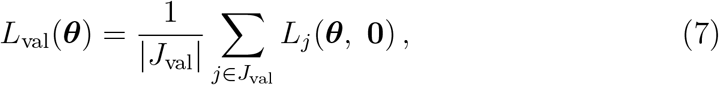

where | *J*_val_ | denotes the cardinality of the validation set.

Parameter updates are performed using the ADAM optimization algorithm [11], which combines adaptive learning rates with momentum to achieve efficient and stable convergence.

We consider two ways of computing gradients:

- **Global step**: The gradient of the total loss *L*_train_ is computed at each optimization iteration. This corresponds to what is commonly referred to as full-batch optimization [28]. We use the term *global step* to avoid confusion with the notion of batch experiment.
- **Minibatch step**: Gradients are computed using a single loss term *L*_*j*_ at a time. In the minibatch setting adopted here, we generate a random permutation of the experiment indexes *J*_train_ at the beginning of each epoch, and the parameters are updated sequentially according to the gradients of the corresponding *L*_*j*_. Therefore, each epoch processes all batch experiments exactly once, while each update reflects the information contained in one batch experiment (see Figure 2).

**Figure 1.**
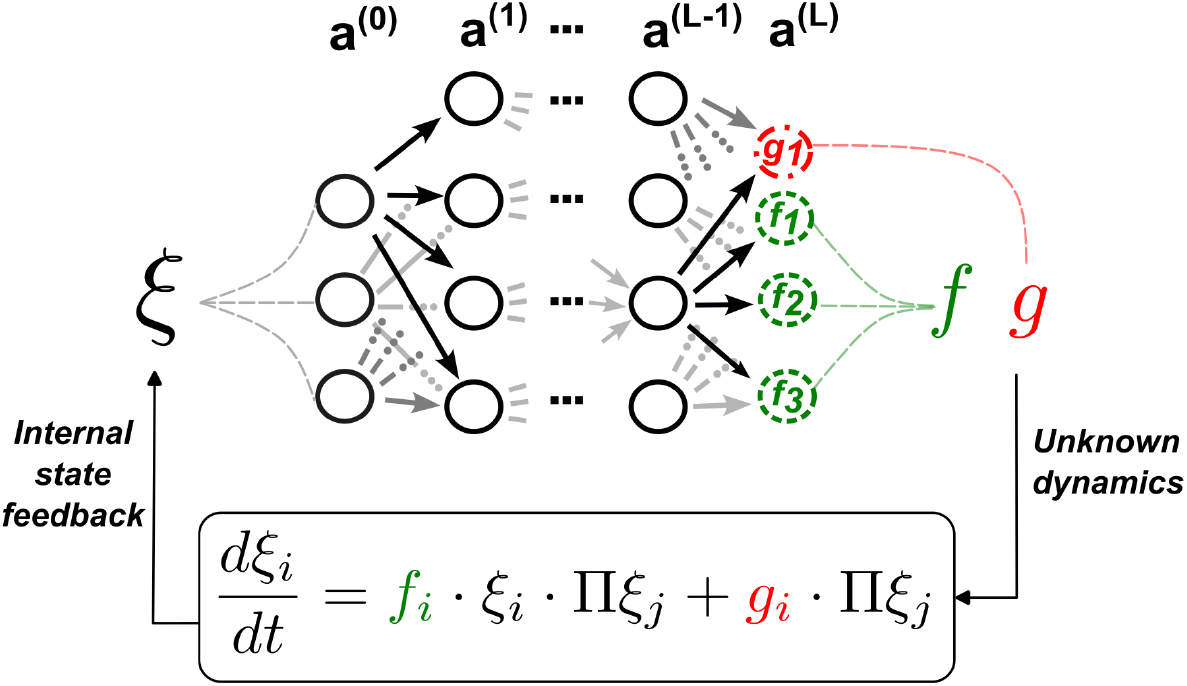
Scheme of the hybrid model. A NN is used to describe the functions *f*_*i*_ and *g*_*i*_, which receive the state vector as input layer. The NN consist of layers *a*^*l*^, *l* ∈ {0, .., *L*}. On the right hand side of the ODE, the functions *g*_*i*_ and *f*_*i*_ are multiplied by state variables. The function *f*_*i*_ is necessarily multiplied by *ξ*_*i*_, which ensures non-negativity of the solutions (see Proposition 1).

**Figure 2.**
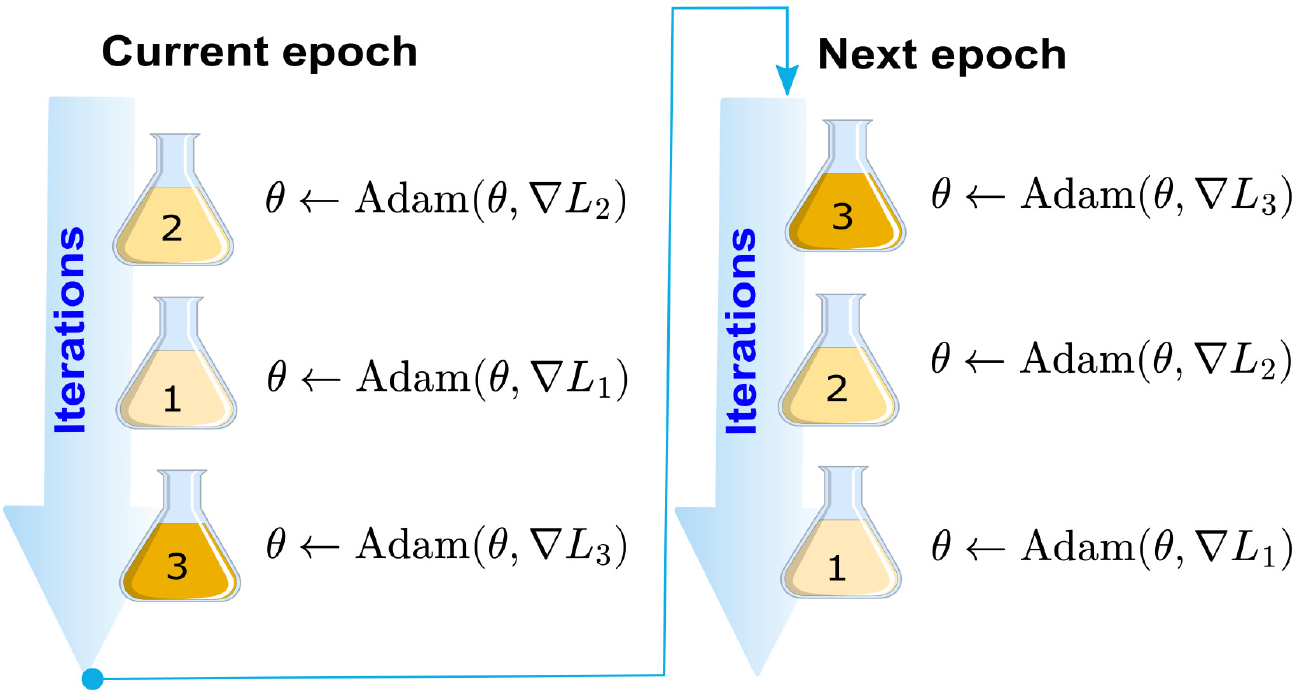
Illustration of the minibatch training strategy considering three batch experiments. In this approach, each batch culture experiment is treated as an individual minibatch, aligning the optimization process with the natural structure of bioprocess data. One epoch corresponds to a full pass through all experiments, with model parameters updated after processing each experiment using the ADAM optimizer. For simplicity, the scheme only illustrates the update of ***θ***, while the experiment-specific initial-condition parameters *α*_*j*_ are omitted.

Because the available data are limited, we apply dropout regularization during training to prevent overfitting [27]. In this scheme, a fixed fraction of neurons in the hidden layers is randomly deactivated at each iteration, forcing the network to learn more robust and distributed representations while maintaining consistent dynamics across each ODE integration.

The neural network training and hybrid ODE simulations are implemented in Python using the JAX library [5] to leverage automatic differentiation and just-in-time (JIT) compilation for efficient computations on TPU devices. The ADAM optimizer from the Optax library was used to update the parameters. All computations, including neural network forward passes and ODE integrations, were JIT-compiled and executed on a TPU v5e-1 device. The hybrid ODE system is solved using the JAX odeint routine with an upper bound on the number of internal steps (mxstep = 2000), introduced to avoid solver stagnation in regions where the dynamics become numerically demanding.

To initialize the weights of the neural network, we adopt a two-step initialization process. First, all weights are drawn i.i.d. from a uniform distribution *U* (−0.1, 0.1). Second, the network parameters are initialized to enforce *f*_*i*_(***ξ, θ***) = 0 and *g*_*i*_(***ξ, θ***) = 0 for all *I* ∈ ℐ, yielding an identically zero vector field. Thus, all state variables remain constant at their initial conditions during initialization, providing a neutral and numerically stable starting point for training. To reduce sensitivity to initialization, the training procedure is repeated ten times with different random initializations in the first step, and the model achieving the lowest validation loss is selected.

## 3 Applications

The purpose of this section is to demonstrate the application of the hybrid modeling framework introduced in Section 2 through two illustrative examples. We outline how to construct hybrid models using Procedure 1, and we compare different training strategies, with particular emphasis on the minibatch approach introduced in Section 2.5. We then assess the predictive performance of the resulting models and examine whether the FNN component exhibits interpretable behavior. A comprehensive architecture search is beyond the scope of this study; instead, we adopt an architecture selected based on initial exploratory tests.

All scripts and data used in this section are publicly available at: https://github.com/carlosmvond/hybrid-modelling-of-batch-bioreactors.

### 3.1 Hybrid model for *E. coli*

We use synthetic data generated with the model proposed by [14], which describes the growth of *E. coli* on glucose. The model accounts for overflow metabolism, meaning that when the glucose uptake rate reaches a threshold, acetate is produced [2]. The model has three state variables: biomass (*X*), glucose (*G*), and acetate (*A*) concentration. To construct the training and validation datasets, we simulate four batch experiments with different initial glucose concentrations: 10, 20, 35, and 50 g/L. Three of these conditions (10, 20, and 50 g/L) are used for training, while the intermediate condition (35 g/L) is reserved for validation. This choice ensures that the validation scenario lies within the range of the training conditions, thus avoiding extrapolation. To further assess the generalization capability of the hybrid model, we additionally generate six independent batch experiments for testing. These test batches cover a broader set of initial glucose concentrations (5, 15, 25, 30, 40, and 45 g/L), allowing us to evaluate model performance across both interpolative and moderately extrapolative scenarios.

To formulate a hybrid model, we follow Procedure 1. Step 1 provides the following seed model:

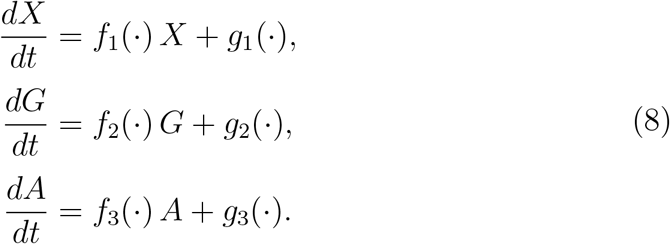

Step 2 identifies the state variables that are zero-invariant, *i*.*e*., variables that cannot increase when their concentration is zero. In this system, biomass *X* and glucose *G* cannot be generated spontaneously in a closed culture; therefore, we remove their corresponding generation terms by setting *g*_1_ ≡ 0 and *g*_2_ ≡ 0. In contrast, acetate *A* can be synthesized by *E. coli* even when *A* = 0; hence, its generation term *g*_3_ is not set to zero.

Step 3 incorporates the mediated interactions among the variables. Since no substrate consumption or acetate formation occurs in the absence of biomass, the variable *X* acts as a mediator that multiplies the corresponding RHSs.

The resulting hybrid model reads:

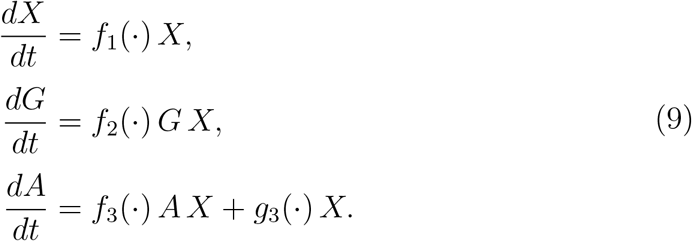

Note that the variable *G* could, in principle, also multiply the RHS of the equation for *X* in (11), assuming that *f*_1_(·) represents the specific growth rate and that growth requires the presence of *G*. However, *f*_1_(·) may take negative values when *G* is depleted, reflecting biomass decay [13]. We also note that by selecting suitable forms for the functions *f*_1_, *f*_2_, *f*_3_, and *g*_3_, model (9) can represent any batch process in which a single metabolite is produced by the microorganism and consumed exclusively through cellular activity.

We evaluate four training configurations using the same FNN architecture, consisting of a single hidden layer with five neurons: (i) minibatch with dropout, (ii) minibatch without dropout, (iii) global training with dropout, and (iv) global training without dropout. All configurations were trained for 2000 epochs using the ADAM optimizer with a learning rate of 0.001 and an initial-condition regularization weight *λ*_*IC*_ = 1. As reported in Table 1, the combination of minibatch training with a dropout rate of 0.01 yields the lowest validation loss. Additionally, the minibatch approach reduces CPU time by approximately a factor of fifteen compared with the global strategy, both with and without dropout. Figure 3 illustrates the quality of the fit obtained using the minibatch+dropout configuration, showing close agreement between model predictions and experimental data across all states. This is further supported by the *R*^2^ values reported in Table 2, which remain consistently between 0.90 and 0.99 for all training and validation batch experiments.

**Table 1.** CPU time, training loss (defined by *L*_train_ in (6)) validation loss (defined by *L*_val_ in (7)) for different training configurations applied to model (9). The updating step refers to the strategy used to update the model parameters (either *batch* or *global*; see Section 2.5). CPU time is the amount of time during which the CPU was actively executing instructions for your process. CPU time is reported after 2000 epochs. Training and validation losses are reported at the epoch where the minimum validation loss was achieved, with the corresponding epoch indicated in parentheses. The configuration yielding the lowest validation loss is highlighted in gray.

| Updating step | Dropout rate | CPU time (s) | Training loss | Validation loss |
| --- | --- | --- | --- | --- |
| Minibatch | 0.1 | 20.75 | 0.118 | 0.062 (epoch 754) |
| Minibatch | 0.01 | 20.81 | 0.031 | 0.013 (epoch 1923) |
| Minibatch | 0 | 21.64 | 0.045 | 0.019 (epoch 481) |
| Global | 0.1 | 313.12 | 0.144 | 0.078 (epoch 961) |
| Global | 0.01 | 317.69 | 0.054 | 0.024 (epoch 1521) |
| Global | 0 | 316.98 | 0.026 | 0.033 (epoch 1969) |

**Table 2.** Determination coefficients (*R*^2^) associated with Figure 3, comparing model simulations with experimental data for biomass (*X*), glucose (*G*), and acetate (*A*). Batches 1–3 were used for training, while Batch 4 served for validation. Batches 5-10 are for testing. The column *G*_0_ indicates the experimental initial glucose concentration.

| Batch | $G_0$ (g/L) | $X$ | $G$ | $A$ |
| --- | --- | --- | --- | --- |
| 1 | 10 | 0.9698 | 0.9847 | 0.9906 |
| 2 | 20 | 0.9927 | 0.9931 | 0.9944 |
| 3 | 35 | 0.9981 | 0.9967 | 0.9937 |
| 4 | 50 | 0.9953 | 0.9930 | 0.9862 |
| 5 | 5 | 0.6590 | 0.4737 | -0.2855 |
| 6 | 15 | 0.9559 | 0.9383 | 0.1548 |
| 7 | 25 | 0.9874 | 0.9959 | 0.9940 |
| 8 | 30 | 0.9825 | 0.9941 | 0.9878 |
| 9 | 40 | 0.9928 | 0.9910 | 0.5948 |
| 10 | 45 | 0.9922 | 0.9901 | 0.6872 |

**Figure 3.**
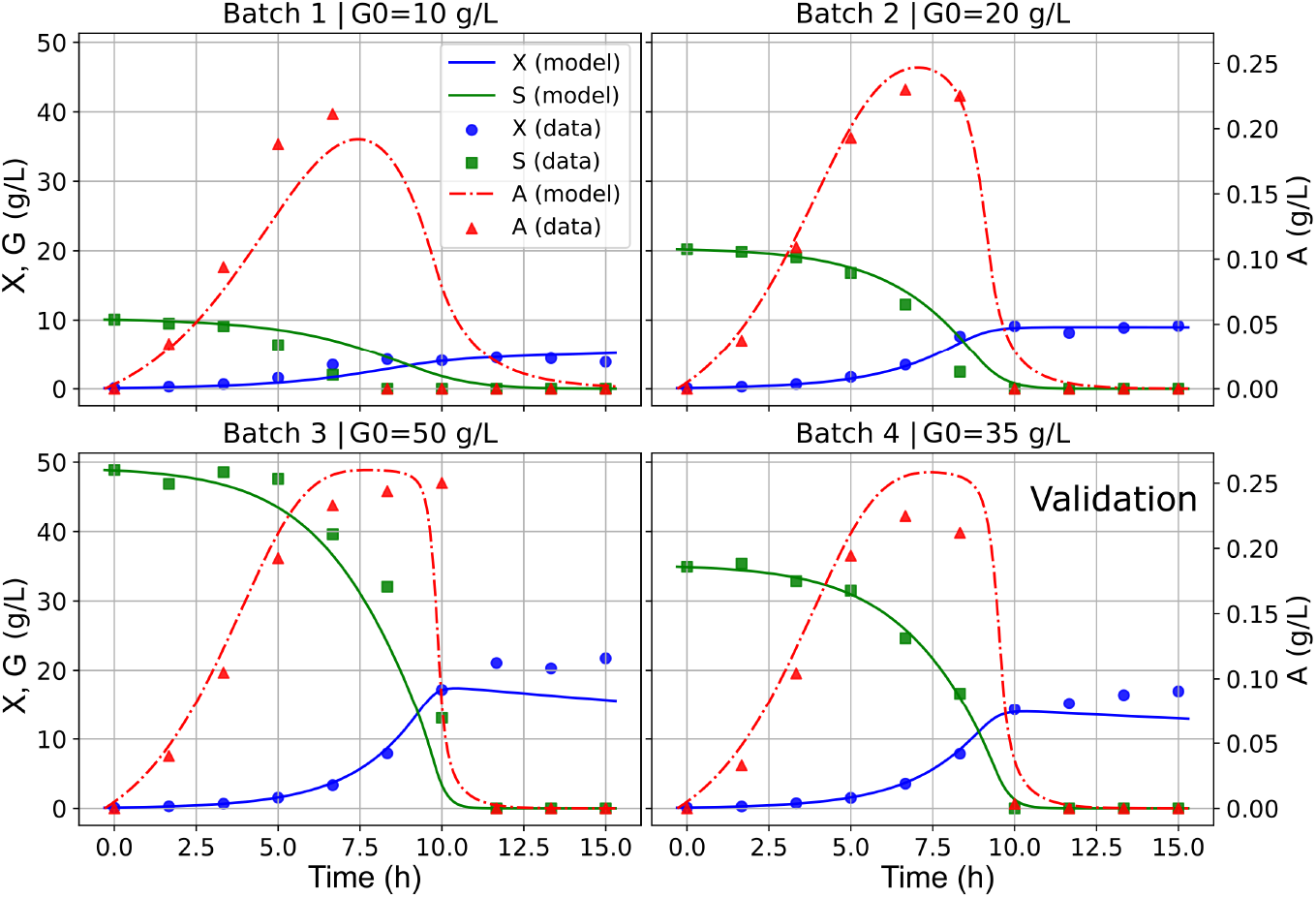
Predicted trajectories generated with model (9) together with the corresponding synthetic data. Solid lines show model predictions, while markers denote the synthetic data points. Biomass (*X*) and glucose (*G*) are plotted on the left axis, and acetate (*A*) on the right axis. Each panel displays the batch number and the initial glucose concentration (*G*_0_). Training was performed using the minibatch strategy with a dropout rate of 0.01, a learning rate of 0.001, and 2000 epochs. Determination coefficients are summarized in Table 2.

To assess whether the FFN provides interpretable information, Figure 4 illustrates the learned relationships between the functions *f*_1_ and *g*_3_ appearing in the hybrid model. Although these functions are highly flexible and could, in principle, take arbitrary shapes, the FNN recovers trends that are biologically meaningful. The function *f*_1_(·) can be interpreted as the specific growth rate, whereas *g*_3_(·) corresponds to the specific acetate production rate. The learned *g*_3_ exhibits a negligible response at low values of *f*_1_ and becomes positive and approximately linear as *f*_1_ increases. This pattern is consistent with the onset of overflow metabolism [2], suggesting that the neural network successfully reconstructs this physiological phenomenon directly from the data.

**Figure 4.**
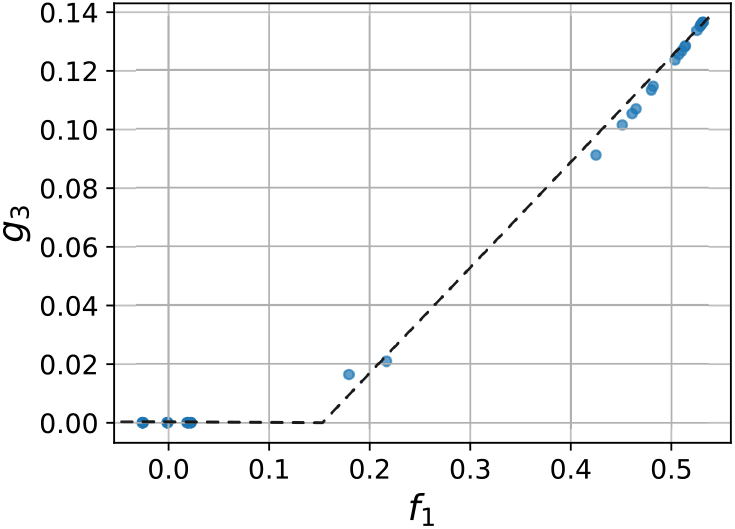
Plot of the relationship between the *f*_1_(·) and *g*_3_(·) learned by model (9) using minibatch training with a dropout rate of 0.01, evaluated at the experimental data.

Regarding the test data set, we observe that five of the six test batches (Batches 6–10) achieve high predictive accuracy for *X* and *G*, with *R*^2^ values between 0.9 and 0.99 for most variables. These conditions lie well within the interpolation region defined by the training experiments. Batches 6, 9, and 10, which are also an interpolation case, show moderate performance with lower *R*^2^ values for acetate. In contrast, Batch 5 (*G*_0_ = 5 g/L) represents true extrapolation beyond the training range and exhibits markedly poorer performance, including negative *R*^2^ values for glucose and acetate. This shows that prediction quality deteriorates when initial conditions fall outside the domain covered during training.

### 3.2 Hybrid model for astaxanthin production

We use experimental data on astaxanthin production by the yeast *Xanthophyllomyces dendrorhous* reported in [12]. The dataset consists of four batch cultures with measurements of biomass (*X*), glucose (*G*), ethanol (*E*), and astaxanthin (*P*) concentrations. Each batch differs in its initial glucose concentration: 25, 50, 75, and 100 g/L. The batches with 25, 50, and 100 g/L are used for training, whereas the batch with 75 g/L is reserved for validation.

Following Procedure 1, Step 1 provides the generic seed formulation:

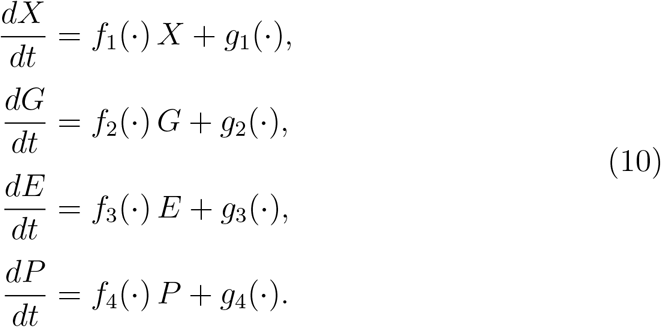

Next, Step 2 identifies the state variables that are zero-invariant. Biomass *X* and glucose *G* cannot be generated spontaneously in a closed culture; therefore, their corresponding generation terms are removed by setting *g*_1_ ≡ 0 and *g*_2_ ≡ 0. In contrast, ethanol *E* and astaxanthin *P* may be produced even when their concentrations are initially zero, provided that biomass is present. Consequently, their generation terms *g*_3_(·) and *g*_4_(·) are retained.

Step 3 incorporates the mediated interactions among the variables. Since substrate consumption cannot occur in the absence of biomass, *X* acts as a mediator and multiplies the corresponding terms on the RHS. The same rationale applies to ethanol: neither its production nor its consumption can proceed without biomass, and therefore *X* multiplies its RHS as well. Astaxanthin (*P*), on the other hand, can only be synthesized in the presence of biomass, yet its rate of change does not vanish when *X* = 0 because degradation may still occur. For this reason, *X* multiplies only the production term *g*_4_(·).

The resulting hybrid model reads:

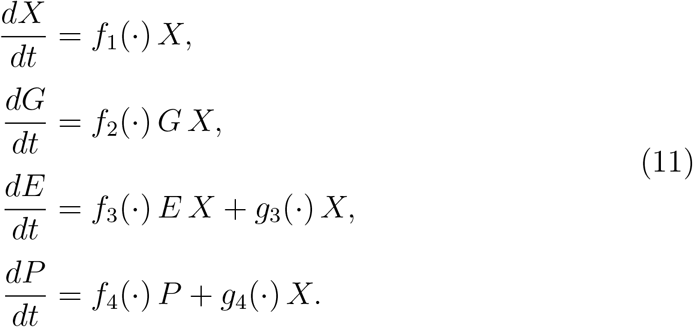

Here, the functions *f*_*i*_(·) and *g*_*i*_(·) denote general state- and input-dependent rates defined within our hybrid modeling framework. This formulation naturally encompasses the model proposed by [12].

#### Remark

The standard model in the literature for describing the production of a metabolite *P* is the Luedeking–Piret model [10], that is:

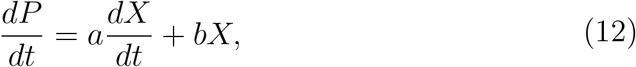

where *a* is the growth-associated production coefficient and *b* is the non-growth-associated production coefficient. However, this model does not ensure the non-negativity of *P*. Indeed, if *P* = 0, nothing rules out the possibility of *dP/dt* being negative, thus leading to negative values of the state variable. Moreover, in the absence of biomass, (12) implies that no change in product concentration occur; however product degradation may occur. Thus, (12) may not be appropriate for describing the dynamics of product concentration.

Similar to the previous section (3.1), to assess both the predictive capability of the proposed hybrid model and the effectiveness of the minibatch training strategy, we compare four training configurations using the same FNN architecture, which consists of four hidden layers with four neurons each: (i) minibatch with dropout, (ii) minibatch without dropout, (iii) global training with dropout, and (iv) global training without dropout. Table 3 shows that the use of minibatch compared to global strategy accelerates more than ten times the method. Regarding the validation loss, the lowest value was reached with minibatch and without dropout.

**Table 3.** CPU time, training loss (defined by *L*_train_ in (6)) validation loss (defined by *L*_val_ in (7)) for different training configurations applied to model (11). The updating step refers to the strategy used to update the model parameters (either *batch* or *global*; see Section 2.5). CPU time is the amount of time during which the CPU was actively executing instructions for your process. CPU time is reported after 2000 epochs. Training and validation losses are reported at the epoch where the minimum validation loss was achieved, with the corresponding epoch indicated in parentheses. The configuration yielding the lowest validation loss is highlighted in gray.

| Updating step | Dropout rate | CPU time (s) | Training loss | Validation loss |
| --- | --- | --- | --- | --- |
| Minibatch | 0.1 | 28.04 | 0.989 | 0.742 (epoch 1829) |
| Minibatch | 0.01 | 29.87 | 0.490 | 0.213 (epoch 983) |
| Minibatch | 0 | 26.95 | 0.296 | 0.125 (epoch 1763) |
| Global | 0.1 | 441.33 | 0.648 | 0.761 (epoch 1267) |
| Global | 0.01 | 454.63 | 0.361 | 0.137 (epoch 1780) |
| Global | 0 | 373.92 | 0.931 | 0.272 (epoch 590) |

Figure 5 shows the results obtained with dropout rate of 0.01 and mini-batch strategy. As summarized in Table 4, the model achieved consistently high determination coefficients in the training batches, with *R*^2^ values ranging from 0.70 to 0.99 for biomass, glucose, and astaxanthin concentrations, indicating that the learned dynamics capture the experimental trends with good accuracy. The validation batch (Batch 4) also shows satisfactory predictive performance, with *R*^2^ values of 0.863, 0.988, 0.949, and 0.982 for biomass, glucose, ethanol, and the product, respectively. These results suggest that the hybrid model generalizes well to unseen conditions despite being trained with a limited number of experiments, demonstrating both robustness and physical consistency in reproducing the coupled growth–production dynamics.

**Table 4.** Determination coefficients (*R*^2^), associated with Figure 5, comparing model simulations with experimental data for biomass (*X*), glucose (*G*), ethanol (*E*), and product (*P*). Batches 1–3 were used for training, while Batch 4 served for validation.

| Batch | $G_0$ (g/L) | $X$ | $G$ | $E$ | $P$ |
| --- | --- | --- | --- | --- | --- |
| 1 | 50 | 0.914 | 0.940 | 0.950 | 0.991 |
| 2 | 100 | 0.702 | 0.943 | 0.682 | 0.935 |
| 3 | 25 | 0.934 | 0.978 | 0.920 | 0.978 |
| 4 | 75 | 0.863 | 0.988 | 0.949 | 0.982 |

**Figure 5.**
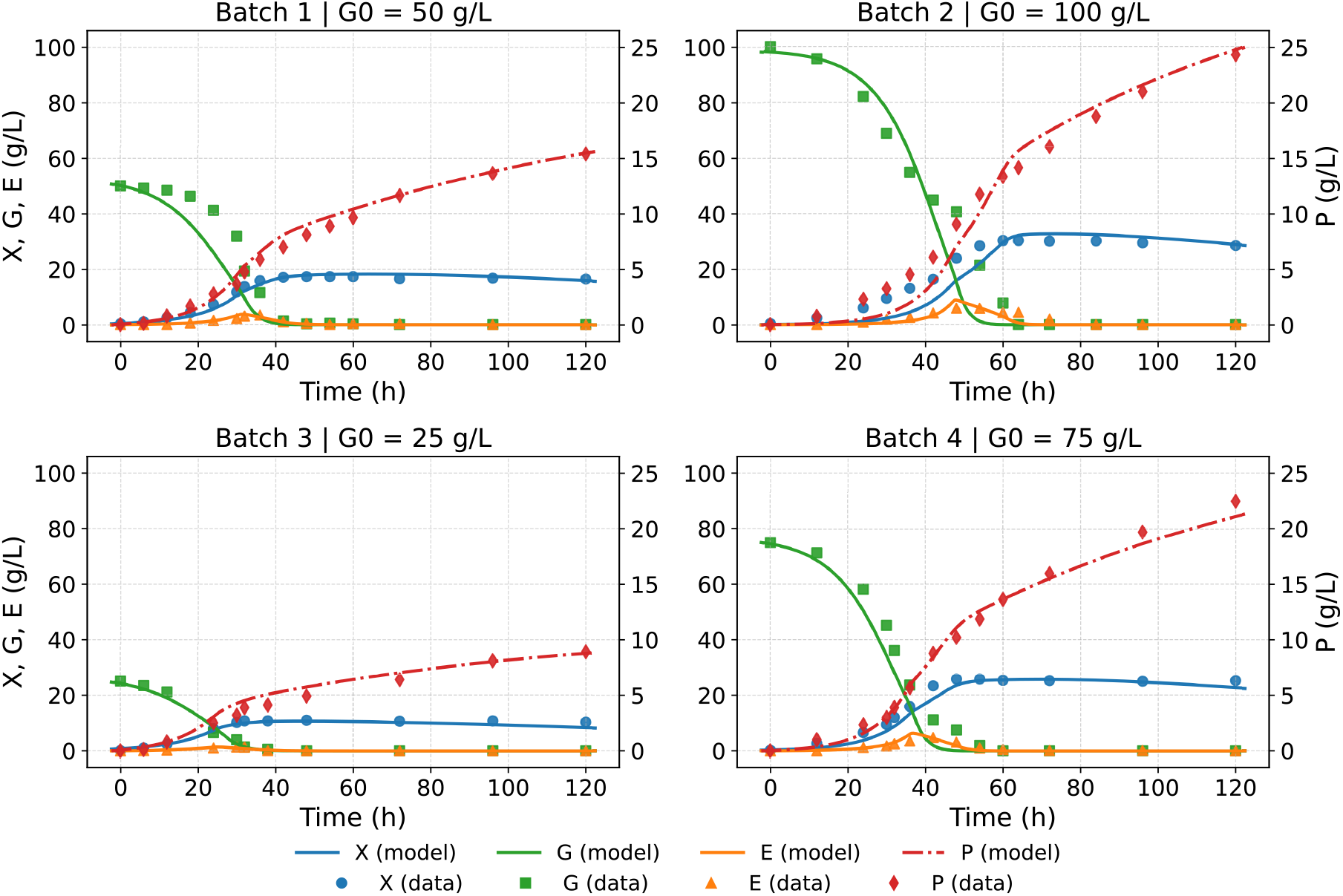
Predicted trajectories generated with model (11) together with the corresponding experimental data. Solid lines show the model predictions for biomass (*X*), glucose (*G*), and ethanol (*E*) on the left axis, and the dash–dot red line shows the product concentration (*P*) on the right axis. Symbols denote experimental measurements. Each panel reports the initial glucose concentration (*G*_0_). Training employed a minibatch scheme without dropout, a learning rate of 0.001, and 2000 epochs. Parameters were taken at the epoch with the lowest validation loss (see Table 3).

Regarding the interpretability of the FNN, Figure 6 reveals clear functional relationships between the learned terms *g*_3_(·), *g*_4_(·), and *f*_1_(·). The function *f*_1_ can be interpreted as the specific growth rate of the yeast, while *g*_3_ and *g*_4_ represent the specific production rates of ethanol and astaxanthin, respectively. As shown in Figure 6, ethanol production (*g*_3_) becomes significant only once *f*_1_ exceeds a threshold value, a behavior consistent with overflow-like metabolic activation and analogous to the trend observed in the previous application (see Figure 4). In contrast, astaxanthin production (*g*_4_) exhibits an increasing trend with respect to *f*_1_, suggesting a direct association between growth and pigment synthesis under the experimental conditions considered. These results highlight that, despite the model data-driven nature, the FNN component is capable of recovering biologically meaningful dependencies directly from the data.

**Figure 6.**
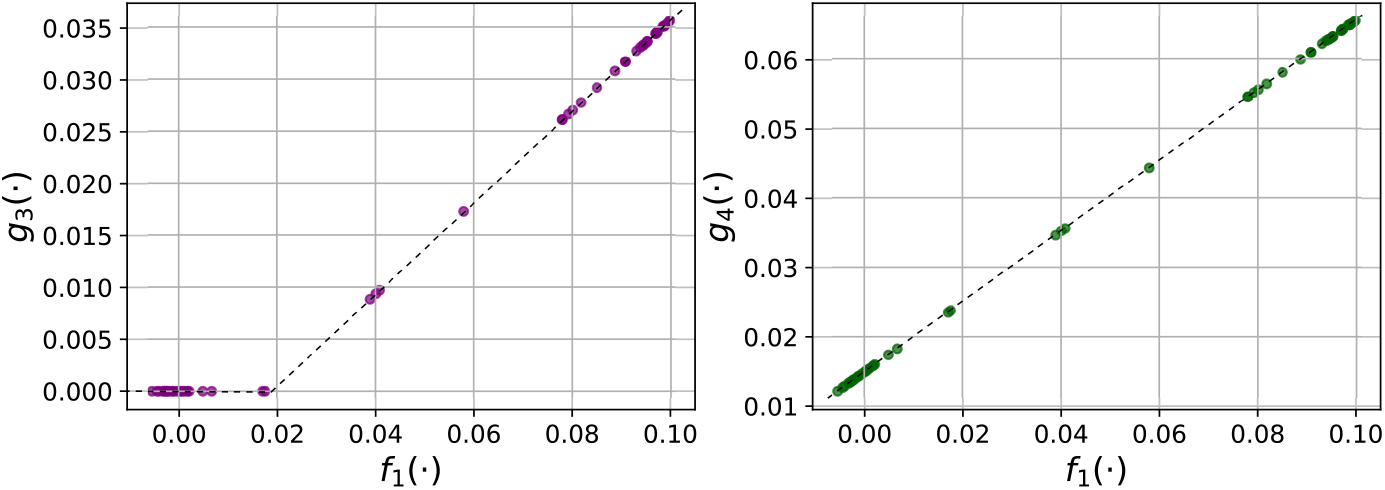
Plot of the relationships between the production rates *g*_3_(·) and *g*_4_(·) and the specific growth rate *f*_1_(·) learned by model (11) using minibatch training without dropout, evaluated at the experimental data.

## 4 Conclusions

We have proposed a framework for constructing bioprocess models using minimal prior knowledge and limited data. The effectiveness of this approach was demonstrated in two case studies: one based on synthetic data and the other on real experimental measurements. In both examples, we found that as few as three batch experiments were sufficient to train a model that reproduced the main dynamical trends of an unseen batch, although prediction accuracy varied across conditions and variables. Furthermore, the neural network components revealed interpretable structures that align with biologically meaningful behaviors, offering not only predictive capability but also valuable insights into the underlying process dynamics.

We proposed a systematic procedure for formulating the mechanistic component of the hybrid model (see Procedure 1). While most of the hybrid model constructions are based on mass balance principles and/or partially known kinetics, our modeling framework relies on minimal prior knowledge and does not introduce any parameters associated with mechanistic knowledge. This design offers two key advantages for parameter estimation. First, it avoids the need to simultaneously estimate parameters from both the mechanistic model and the machine learning component, a major challenge in hybrid modeling [25]. Second, we incorporate prior knowledge that is certain and well established. This point is crucial since including incorrect or uncertain information can degrade model performance and lead to ill-conditioned estimation problems [23]. Together, these features confer a high degree of flexibility to the proposed framework, making it well suited for a broad range of bioprocess scenarios, including those requiring adaptive or incrementally updated models [6].

We introduced a minibatch training strategy inspired by machine-learning practices but tailored to the structure of batch bioprocess data. Unlike the conventional minibatch approach, in which randomly selected data points are used to compute stochastic gradient updates, our method assigns each minibatch to a complete batch experiment. This design exploits the natural organization of bioprocess data and avoids repeated ODE integrations with heterogeneous initial conditions within a single update. In Application 1, the minibatch scheme achieved a training loss comparable to that of the global-step method but substantially reduced CPU time. In Application 2, it not only accelerated training but also improved model fitting. These results demonstrate that adapting classical optimization algorithms to domain-specific data structures can yield both computational and predictive advantages.

Finally, the proposed framework offers new possibilities for developing interpretable models. Whereas recent efforts rely on symbolic regression [9, 24], our formulation enables interpretable structures to be extracted directly from the trained hybrid model, opening a complementary path for identifying underlying dynamics.

Overall, this work introduces a flexible and interpretable hybrid modeling strategy for bioprocesses. By avoiding strong mechanistic assumptions and exploiting the structure of batch experimental data, the framework provides a practical and efficient alternative to traditional hybrid models, contributing to the development of robust modeling tools for early-stage bioprocess research.

## Acknowledgments

This work was supported by the National Research and Development Agency (ANID) through FONDECYT Grant No. 11251726, and by the Pontificia Universidad Católica de Valparaíso through Grant No. 039.770/2025. This work was partially supported by ITMO Cancer of Aviesan as part of the 2021–2030 Cancer Control Strategy (MIC proposal Cellema, grant 22CM045-00), via funds administered by Inserm to Jérémie Roux, who served as the postdoctoral supervisor of Marielle Péré.

## Data availability statement

The datasets used for model training in this study were obtained from previously published works. Specifically, the synthetic dataset for *E. coli* growth was generated using the model proposed by Mauri et al. [14], while the experimental data for astaxanthin production by *X. dendrorhous* were taken from Liu and Wu [12].

The scripts used for training the models in Section 3 are available in https://github.com/carlosmvond/hybrid-modelling-of-batch-bioreactors.

